# Functional analysis of a biosynthetic cluster essential for production of 4-formylaminooxyvinylglycine, a germination-arrest factor from *Pseudomonas fluorescens* WH6

**DOI:** 10.1101/080572

**Authors:** Rachel A. Okrent, Kristin M. Trippe, Maciej Maselko, Viola Manning

## Abstract

Rhizosphere-associated *Pseudomonas fluorescens* WH6 produces the germination-arrest factor, 4-formylaminooxyvinylglycine (FVG). FVG has previously been shown to both arrest the germination of weedy grasses and to inhibit the growth of the bacterial plant pathogen *Erwinia amylovora.* Very little is known about the mechanism by which FVG is produced. Although a previous study identified a region of the genome that may be involved in FVG biosynthesis, it has not yet been determined which genes within that region are sufficient and necessary for FVG production. In the current study, we explored the role of each of the putative genes encoded in that region by constructing deletion mutations. Mutant strains were assayed for their ability to produce FVG with a combination of biological assays and thin-layer chromatographic analyses. This work defined the core FVG biosynthetic gene cluster and revealed several interesting characteristics of FVG production. We determined that FVG biosynthesis requires two small open reading frames of less than 150 nucleotides and that multiple transporters have overlapping but distinct functionality. In addition, two genes in the center of the biosynthetic gene cluster are not required for FVG production, suggesting that additional products may be produced from the cluster. Transcriptional analysis indicated that at least three active promoters play a role in the expression of genes within this cluster. The results of this study enrich our knowledge regarding the diversity of mechanisms by which bacteria produce non-proteinogenic amino acids like vinylglycines.

## INTRODUCTION

*Pseudomonas fluorescens* WH6, a bacterial strain originally isolated from the rhizosphere of wheat (Elliott *et al.*, 1998), has been investigated in our laboratory for its ability to produce a secondary metabolite with selective herbicidal and antimicrobial properties. WH6 culture filtrate arrests the germination of a number of weedy grass species, including annual bluegrass (*Poa annua* L.) (Banowetz *et al.*, 2008; Banowetz *et al.*, 2009). WH6 filtrate also inhibits the growth of the bacterial plant pathogen *Erwinia amylovora* (Halgren *et al.*, 2011), the causal agent of fireblight in orchard crops. Based on the biological effects of the culture filtrates on grass seed germination, the active compound was originally termed a germination-arrest factor (GAF). The molecule responsible for these activities was subsequently purified and identified as 2-amino-4-formylaminooxy-3-butenoic acid, also known as 4-formylaminooxyvinylglycine (FVG) (Fig. 1) (Armstrong *et al.*, 2009; McPhail *et al.*, 2010).

**Fig 1.**
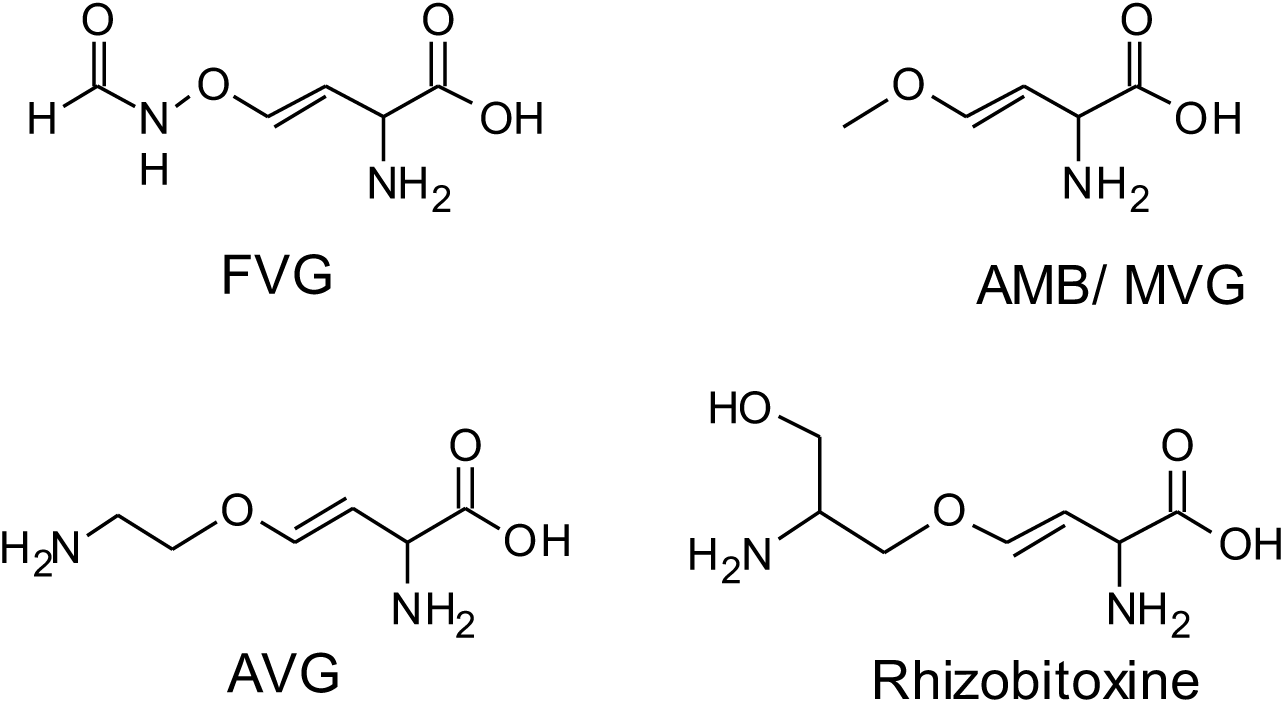
Structures of the GAF molecule 4-formylaminooxyvinylglycine (FVG) and other oxyvinylglycines including 4-methoxyvinylglycine (AMB/MVG) from *P. aeruginosa*, aminoethoxyvinylglycine (AVG) from *Streptomyces* sp., and rhizobitoxine from *Bradyrhizobium elkanii* and *Burkholderia andropogonis.*

FVG is an oxyvinylglycine, a class of non-proteinogenic amino acids produced and excreted by plant-and soil-borne bacteria. Several oxyvinylglycines have been investigated for their role in plant-microbe and microbe-microbe interactions, including aminoethoxyvinylglycine (AVG) from an isolate of *Streptomyces sp.* (Pruess *et al.*, 1974), 4-methoxyvinylglycine (MVG/AMB) from *P. aeruginosa* (Scannell *et al.*, 1975), and rhizobitoxine from *Bradyrhizobium elkanii* (Owens *et al.*, 1972) and *Burkholderia andropogonis* (Mitchell & Frey, 1988) (Fig. 1). The biological activity of oxyvinylglycines is based on their ability to inhibit enzymes that require pyridoxal phosphate (PLP) as a co-factor (Berkowitz *et al.*, 2006). PLP-requiring enzymes are required for important cellular processes including amino acid biosynthesis, nitrogen metabolism (Percudani & Peracchi, 2003), and ethylene biosynthesis (Yu *et al.*, 1979). Because oxyvinylglycines target a broad spectrum of critical enzymes, they can have economically important agricultural applications, including plant growth regulation (Çetinbaş *et al.*, 2012) and weed control (Lee *et al.*, 2013).

Predicting the biosynthetic pathway of FVG using comparative genomics or other tools is difficult because the known biosynthetic pathways of oxyvinylglycines vary considerably (Cuadrado *et al.*, 2004; Rojas Murcia *et al.*, 2015; Yasuta *et al.*, 2001) and because FVG has an unusual aminooxy linkage (Fig. 1). Therefore, determining which genes are required for production of FVG can inform our understanding of its biosynthetic pathway. We previously identified several loci required for production of FVG by screening Tn*5* mutant libraries of the sequenced strain *P. fluorescens* WH6 (Halgren *et al.*, 2013; Kimbrel *et al.*, 2010). Tn*5* insertions in seven regions of the chromosome led to the loss of FVG production. Of these, two occurred in regions that appeared to be specific to FVG production. One of the Tn*5* insertions disrupted a gene encoding the anti-sigma factor PrtR. The role of PrtR was recently investigated and it was found to modulate negative regulation of FVG production through the ECF-type sigma factor PrtI (Okrent *et al.*, 2014). A second 13-kb region was disrupted by multiple Tn*5* insertions that led to the loss of FVG production. This chromosomal region consisted of 12 ORFs that encode putative regulatory proteins, biosynthetic enzymes and transporters (Halgren *et al.*, 2013) (Fig. 2).

**Fig 2.**
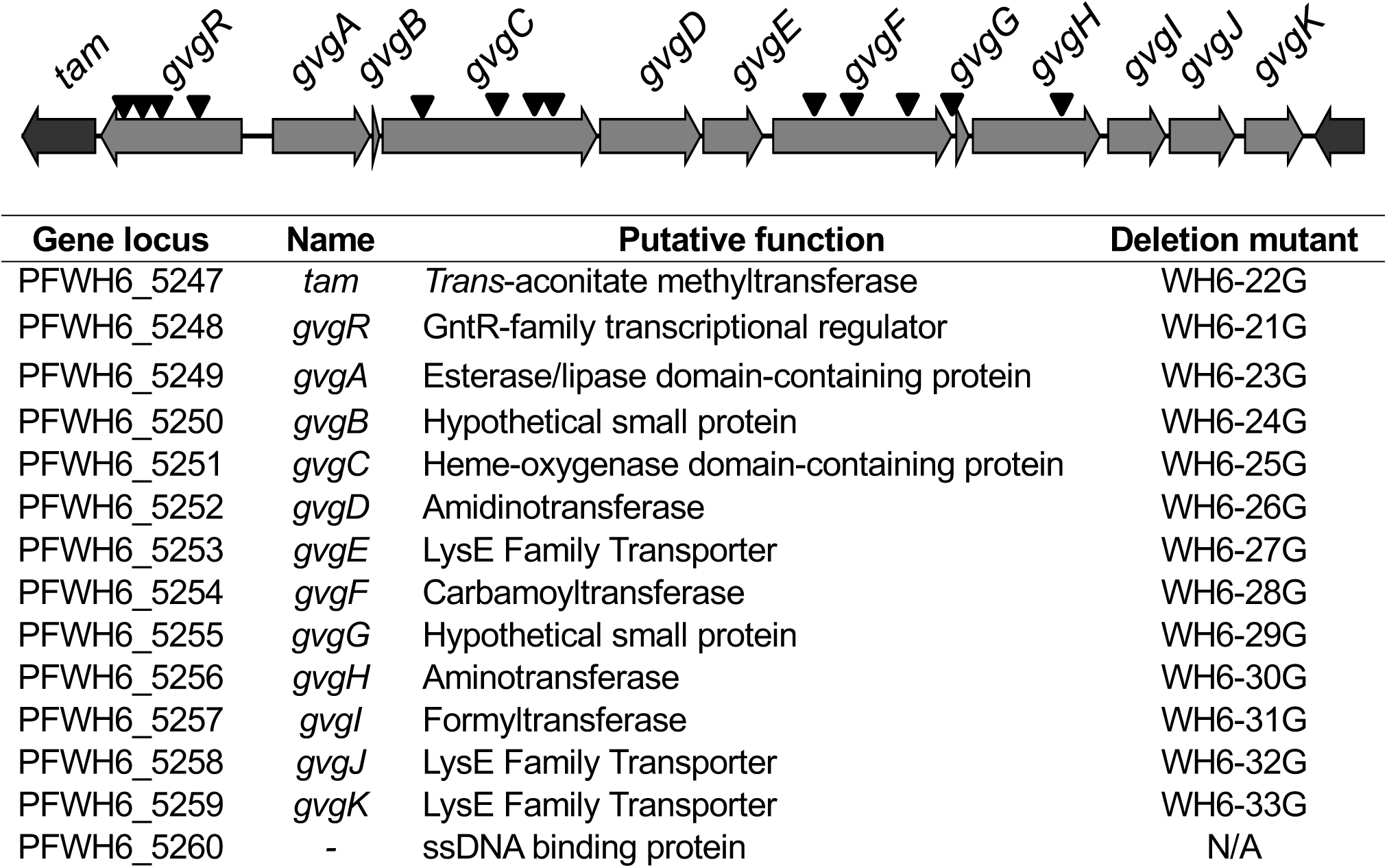
Genetic structure of the *gvg* biosynthetic gene cluster and adjacent genes. The sites of the Tn*5* insertions described in Halgren *et al.* (2013) are indicated with inverted triangles.

In the current study, we characterize this chromosomal region, referred to here as the GAF vinylglycine (*gvg*) cluster. Specifically, we determined which of these genes are necessary for production of FVG through site-directed mutagenesis and complementation. Using this approach, we confirmed the importance of the four genes identified in the initial Tn*5* screens (Halgren *et al.*, 2013; Kimbrel *et al.*, 2010) and identified six additional genes that are involved in the production of FVG. We also investigated the transcriptional organization of the cluster and determined that, in addition to a large polycistronic transcript extending the length of the cluster, there are also several smaller transcripts. This work reveals several interesting aspects of the *gvg* biosynthetic gene cluster, including two small ORFs that are required for production of FVG and multiple genes encoding LysE family transporters with distinct but overlapping functionality. The presence of two genes within the *gvg* cluster that are not required for FVG production suggests that additional products may be produced from the gene cluster.

## METHODS

### Bacterial strains and plasmids

Bacterial strains are listed in **Table S1** and plasmids in **Table S2**. The origin and characterization of *P. fluorescens* strain WH6 (NRRL #B-30485) from wheat rhizosphere was described previously (Banowetz *et al.*, 2008; Elliott *et al.*, 1998).

### DNA manipulation

DNA was isolated from bacteria using the ZR Fungal/Bacterial DNA Kit (Zymo Research) or the PowerLyzer Ultraclean Microbial DNA Isolation Kit (Mo Bio). Purity and concentration were determined using a Nanodrop ND1000 (Thermo Scientific) or Eon Plate Reader with Take3 attachment (Biotek). PCR was performed using high fidelity polymerases when required [i.e. Phusion and Q5 (New England Biolabs (NEB)), Accuprime (Invitrogen)] or OneTaq (NEB) for standard PCR. PCR products were gel purified using standard kits. Restriction enzymes, T4 DNA ligase, and HiFi Assembly reagent were purchased from NEB. Sequencing was performed using an ABI 3730 capillary DNA sequence system (Applied Biosystems) by the Center for Genome Research and Biocomputing Core Facilities (Oregon State University).

### Construction of in-frame deletion mutants

All primers used for PCR amplification in this study are listed in **Table S3**. Site-directed mutagenesis was performed generally as described in (Okrent *et al.*, 2014). However, a scar-less method that omits the insertion of the kanamycin resistance-FLP recognition target sites (*kan*-FRT) cassette was used in some deletions, allowing for the subsequent deletion of additional genes. The pEX-18Km vector was designed for this purpose, with a kanamycin resistance gene (*kan)* from pKD13 (Datsenko & Wanner, 2000) replacing the tetracycline resistance gene (*tet)* of pEX18-Tc. The pEX-18Km vector increased transformation efficiency relative to pEX-18Tc in the scarless method (data not shown). Primers for overlap extension PCR are described in **Table S4** and primers and restriction recognition sites for use in the generation of the FRT-*kan*-FRT fragment are described in **Table S5**.

Plasmids were mobilized into recipients through triparental mating using pRK2013 (Figurski & Helinski, 1979) as the mobilizing helper plasmid. Transconjugants that had undergone a recombination event were selected on 925 minimal medium agar (Halgren *et al.*, 2011) containing 50 µg ampicillin ml-1 and 50 µg kanamycin ml-1. Transconjugants were transferred to plates containing 5% sucrose to counter-select for double recombinants that contain only the mutant allele. When required, eviction of *kan* was mediated by pBH474 (House *et al.*, 2004), which encodes the FLP site-specific recombinase, and confirmed by replica plating on agar plates with and without kanamycin (House *et al.*, 2004). Mutants were identified by colony PCR using primers described in **Table S6**. Each deletion mutant was further confirmed by amplifying the region encompassing the flanking regions, ligating this region into the PCR cloning vector pJET1.2/blunt (CloneJET PCR cloning kit, Thermo Scientific), and sequencing the insert **(Table S6)**.

### Genetic complementation

The deletion mutations in *P. fluorescens* WH6 that resulted in a null or altered FVG phenotype were complemented by inserting the corresponding gene under the control of the constitutive *lac* promoter in the expression vector pBBR1EVM (Okrent *et al.*, 2014) (**Table S7**). The resulting constructs were transferred from *E. coli* into the appropriate WH6 mutant strains by triparental mating as above. Transconjugants expressing resistance to gentamicin (from the plasmid) and ampicillin (from WH6) were selected. Correct transfer of plasmids was confirmed by isolation of the plasmids and digestion with the appropriate restriction enzymes. In some cases, complementation with a native promoter or inducible promoter construct was warranted. Native promoter constructs included sequence upstream of the gene(s) of interest and were constructed in the expression vector pBBR1MCS-5 (Kovach *et al.*, 1995). Inducible promoter constructs were under the control of the *araC*-pBAD system in pJN105 (Newman & Fuqua, 1999).

### Bioassays for FVG activity

*P. fluorescens* strains were inoculated into modified Pseudomonas Minimal Salts Medium (PMS Medium), cultured at 28 °C for 7 days, and harvested as described previously (Banowetz *et al.*, 2008). Filtrates were made from duplicate or triplicate clones of each deletion mutant or complemented mutant strain and subsequently assayed to ensure reproducibility. The bioassay for germination arrest activity in the bacterial culture filtrates was performed using the standard germination arrest bioassay protocol and scoring system described in (Banowetz *et al.*, 2008) for annual bluegrass seeds. The agar diffusion bioassays for anti-microbial activity against *Erwinia amylovora* were performed as described by Halgren *et al.* (2011) with the resulting zones of inhibition measured on triplicate plates using the imaging software Able Image Analyser (Mu Labs).

### TLC analyses

Samples were prepared for TLC analysis by extracting the solids from dried bacterial culture filtrates as described previously (Armstrong *et al.*, 2009; Halgren *et al.*, 2011; Okrent *et al.*, 2014). TLC analysis was performed on Silica Gel GHL and microcrystalline cellulose TLC plates (250 µm thick; Analtech) with ninhydrin-staining as detailed in (Armstrong *et al.*, 2009).

### Construction of the double transporter mutant and test for lethality

The Δ*gvgJK* deletion mutation was first attempted in a wild-type *P. fluorescens* WH6 background using the suicide plasmid pEXW34. As this attempt was unsuccessful, the Δ*gvgJK* mutation was then constructed in the WH6-25G (Δ*gvgC*) background to avoid the potential for the accumulation of toxic products in the cell. The pEXW34 plasmid was mobilized into WH6-25G through triparental mating, resulting in WH6-34G, a triple mutant in *gvgC*, *gvgJ*, and *gvgK*.

The *gvgC* gene was cloned into the broad-host range vector pJN105 (Newman & Fuqua, 1999) under the control of the pBAD arabinose-inducible promoter. This construct, pIJW5251, was mobilized first into WH6-25G (Δ*gvgC*) and assayed for function of *gvgC* in the agar diffusion assay and then subsequently mobilized into the triple mutant WH6-34G. The wild-type, mutant, and complemented mutant strains were grown for 24 h in a modified PMS medium containing glycerol instead of glucose as the carbon source in the presence or absence of 100 mM L-arabinose (Ara). As an indicator of growth, the OD_600_ for triplicate cultures was measured with an Eon Plate Reader (Biotek) with a path length 0.5 cm.

### cDNA synthesis and RT-PCR

RNA was extracted from WH6 bacterial cultures grown to mid-exponential phase in PMS medium and stopped in RNA Protect Bacterial Reagent (Qiagen). The total RNA was extracted using an RNeasy Mini kit (Qiagen) followed by the removal of genomic DNA with Turbo DNase treatment (Ambion). cDNA was synthesized using the SuperScript III First-Strand Synthesis System (Invitrogen) under standard conditions with random hexamer primers. A reaction without Reverse Transcriptase (RT) using the same RNA was performed for the no-RT control sample. To detect transcripts across genes, cDNA from three pooled RNA samples was amplified in PCR reactions with overlapping primer pairs **(Tables S8),** using cDNA as a template. Amplification reactions with no RT and gDNA controls were also included.

### 5′ RACE

Rapid amplification of cDNA ends (RACE) was carried out with the GeneRacer^TM^ Kit (Invitrogen) to determine the transcriptional start sites for *gvgF* and *gvgH*. RNA was purified as above and enriched for full-length transcripts by treatment with Terminator (Epicentre). Full length transcripts with triphosphate ends were subsequently converted to monophosphate 5′ ends with tobacco acid pyrophosphatase. The GeneRacer^TM^ RNA Oligo was then ligated to the treated monophosphate 5′ cDNA ends along with untreated control samples. The WH6 mRNA was reverse transcribed with random hexamers and the resulting cDNA amplified using the GeneRacer^TM^ forward primer and reverse primers specific to each of the two genes examined in nested reactions (**Table S9**). The major PCR products were gel purified and cloned into the pCR4-TOPO™ cloning vector (Invitrogen). Colony PCR was performed on at least ten clones from each PCR band to estimate insert size and the largest inserts from each set were sequenced to determine the location of 5′ ends.

### Construction of *lacZ* reporter fusions and assay for β-galactosidase activity

The putative promoter regions upstream of *gvgF* (493 nt) and *gvgH* (705 nt) were cloned into the mini-Tn*7* vector pXY2 for constructing fusions with *lacZ* to test for promoter activity (Liu *et al.*, 2014). The *lac* promoter was also amplified from pBBR1EVM and cloned into pXY2 as a positive control. Primers are described in **Table S7**. The promoter-*lacZ* fusions were transferred into the conserved attTn*7* site of the wild-type WH6 genome by introduction of the pXY2-based plasmids and pTNS3 that encodes the site-specific Tn7 transposition pathway (Choi *et al.*, 2008) via electroporation as described by Choi *et al.* (2006). Ten colonies from each transformation were patched on to LB plates supplemented with 50 µg ampicillin ml-1, 15 µg gentamicin ml-1 and 40 µg X-Gal ml-1. Plates were incubated at 28 ºC for 48 hours for visualization of β-galactosidase activity. Genomic DNA was extracted from three of the patched isolated per transformation and screened for the genomic insertion with the PCR primers lacZ6 (Liu *et al.*, 2014) and VM029 (**Table S3**).

### Bioinformatics analyses

The locations of putative transmembrane (TM) domains were determined using the web interface of TMHMM v.2.0 from the Center for Biological Sequence Analysis at the Technical University of Denmark (http://www.cbs.dtu.dk/services/TMHMM/). Protein secondary sequence predictions were performed using the PSIPRED Protein Sequence Analysis Workbench from the Bloomsbury Centre for Bioinformatics at University College London (http://bioinf.cs.ucl.ac.uk/psipred/). Promoter sequences were predicted using BPROM from Softberry (http://linux1.softberry.com/) (Solovyev & Salamov, 2011). Rho-independent transcriptional termination sequences were probed using ARNold from the Université Paris-Sud (http://rna.igmors.u-psud.fr/toolbox/arnold/index.php/) (Naville *et al.*, 2011). Potential signal sequences were analyzed using SignalP 4.1 from the Technical University of Denmark (http://www.cbs.dtu.dk/services/SignalP/) (Petersen *et al.*, 2011).

The *P. fluorescens* WH6 GvgF sequence was aligned with characterized O-carbamoyltransferase (Parthier *et al.*, 2012) sequences using MUSCLE v.3.8.425 implemented in CLC Main Workbench. The pairwise identity between the sequences was calculated based on the alignment using the pairwise comparisons tools in the same program. Co-conservation of genes was investigated using the Conserved Neighborhood tool of the Integrated Microbial Genomics platform from the U.S. Department of Energy (http://img.jgi.doe.gov/) (Markowitz *et al.*, 2014).

## RESULTS

### General features of the *gvg* biosynthetic gene cluster

There are 12 predicted open reading frames encoded within the 13-kb chromosomal region that range in length between 84 nt and 2211 nt. Intergenic regions vary from 13 nt to 307 nt. A predicted GntR-family regulator, *gvgR,* is encoded on the opposite strand in relation to the other 11 ORFs (Fig. 2). The *gvg* cluster is flanked by genes encoding a putative *trans*-aconitate methyltransferase and a single-stranded DNA binding protein (Fig. 2). In the case of several genes *(gvgB, gvgC* and *gvgJ)*, the start sites used for constructing deletion mutations and complements were different from the start sites annotated in the published genome (Kimbrel *et al.*, 2010) **(Fig. S1).**

### Effect of deletion mutations and complements on FVG production

FVG activity is measured using a germination arrest assay with annual bluegrass seeds (Banowetz *et al.*, 2008; Banowetz *et al.*, 2009) and an agar diffusion assay with *E. amylovora* (Halgren *et al.*, 2011). Additionally, FVG reacts with ninhydrin to produce either a pink band on silica TLC plates or a purple band on cellulose TLC plates, each at a characteristic R_f_ value (Armstrong *et al.*, 2009). The absence of one or more of these elements in sterile or extracted culture filtrates is indicative of a null FVG phenotype. In an earlier study, Tn*5* insertions in four of the *gvg* genes resulted in the loss of FVG production (Halgren *et al.*, 2011; Kimbrel *et al.*, 2010). These genes, *gvgR, gvgC, gvgF*, and *gvgH*, are predicted to encode a GntR-family transcriptional regulator, a heme-oxygenase domain-containing protein, a carbamoyltransferase (CTase), and an aminotransferase, respectively (Fig. 2). However, Tn5 insertions were not recovered in the other genes within the 13-kb region. Therefore, we used site-directed mutagenesis to determine whether these additional genes are also required for FVG production and to confirm that the phenotype reported in earlier Tn*5* studies were not due to polar effects.

Collectively, site-directed deletion of seven genes in the *gvg* cluster resulted in an obviously null FVG phenotype (Fig. 3, Fig. 4, Table 1). In addition to the four genes originally identified in the screen of Tn*5* insertion mutants (Halgren *et al.*, 2011; Kimbrel *et al.*, 2010), a null FVG phenotype was also observed in site-directed deletions of *gvgA*, encoding an esterase/lipase, and two small ORFs, *gvgB* and *gvgG.* Complementation of these seven deletions with the corresponding wild-type allele *in trans* restored the wild-type FVG phenotype (Fig. 3, Table 1). The mutation in the gene encoding the GntR transcriptional regulator (*gvgR)* could not be successfully complemented with *gvgR* under the control of a constitutive promoter, however it was complemented with a construct containing *gvgR*, *gvgA* and the intergenic region containing the putative promoters of both genes (Fig. 3, Table 1). The mutation in the esterase/lipase *(gvgA)* was complemented with a construct containing the *gvgA-gvgB* sequence under the control of a constitutive promoter. In contrast, mutations in two genes within the cluster, *gvgD* encoding an amidinotransferase and *gvgE* encoding a LysE family transporter, did not affect FVG production (Fig. 3, Fig. 4, Table 1). Mutation of the gene located downstream of *gvgR*, *tam,* did not alter the FVG phenotype (data not shown) and was thus determined to not be part of the *gvg* cluster.

**Fig 3.**
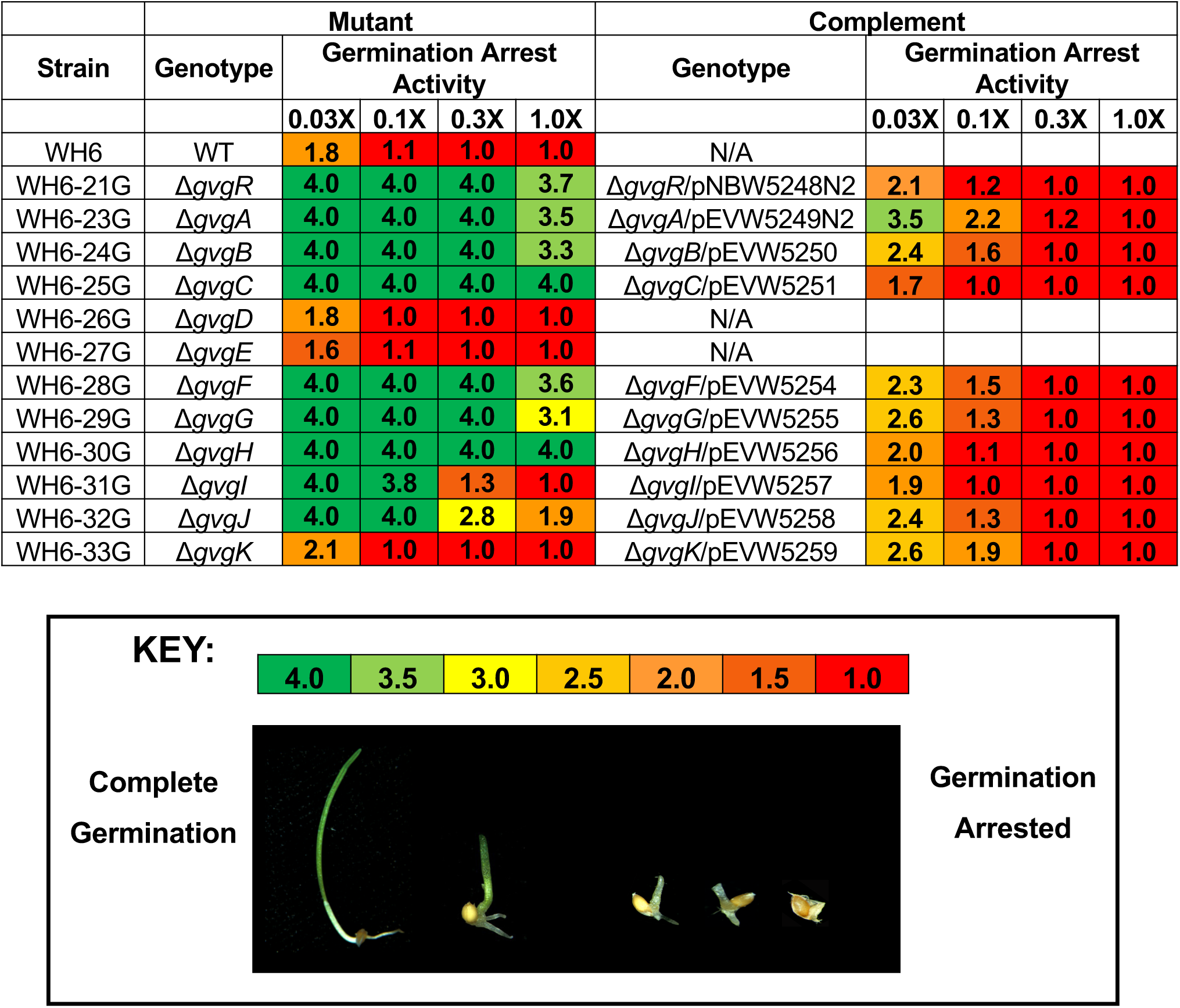
Germination arrest activity of filtrates from *P. fluorescens* WH6 mutants and complements. Annual bluegrass seeds were challenged with a dilution series of culture filtrate from the indicated WH6 mutant strain or complemented mutant strain. In this semi-quantitative assay, a score of 4.0 indicated normal germination and a score of 1.0 indicates that the germination is arrested. The color gradient is from dark green (4.0) to red (1.0) with gradations of color to the nearest 0.5. Corresponding data including standard errors are shown in Tables S10 and S11.

**Fig 4.**
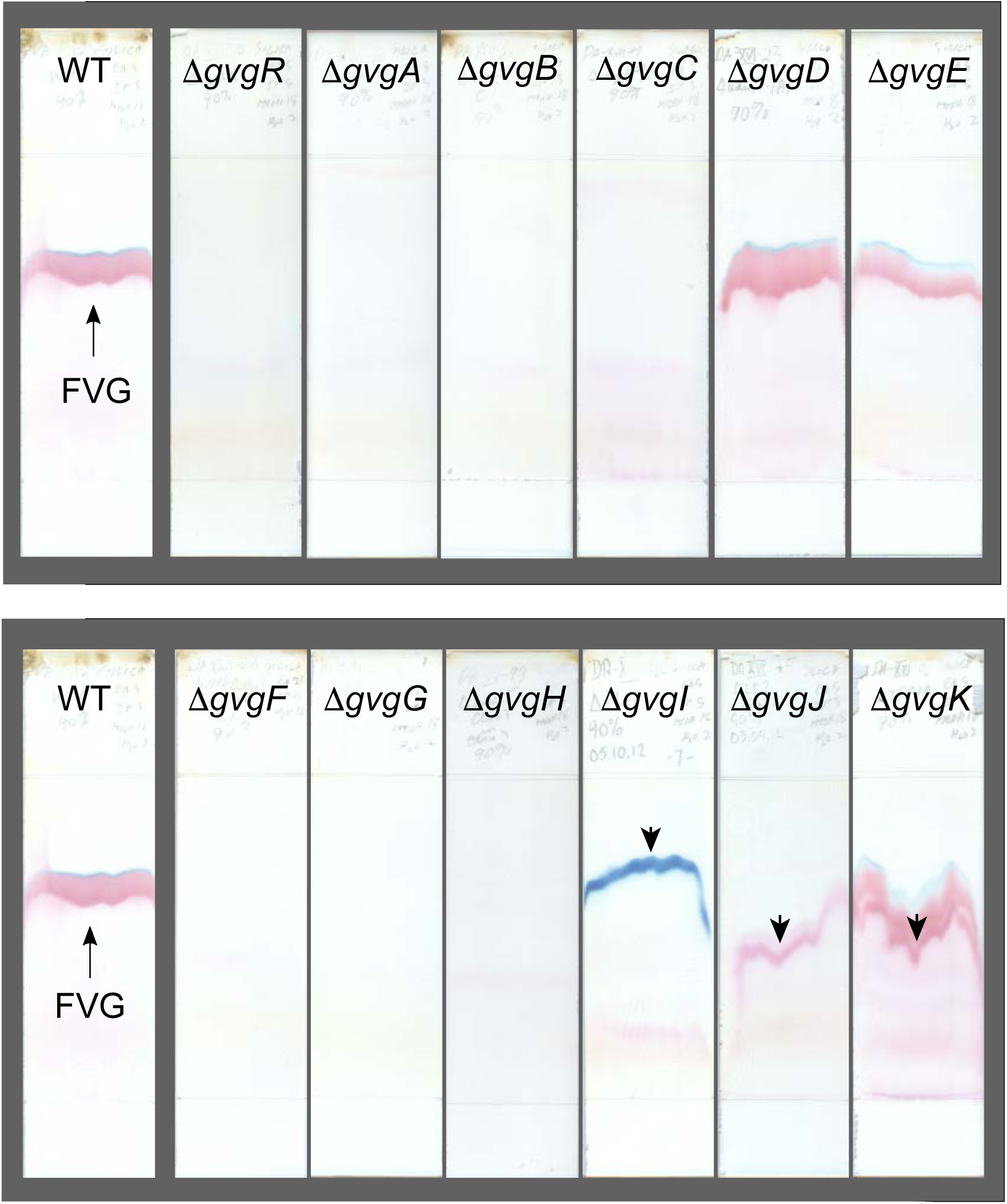
TLC analyses of culture filtrates from *P. fluorescens* WH6 wild-type and mutant strains. Ethanol extracts of evaporated culture filtrates were separated on silica TLC plates and stained with ninhydrin. Downward-pointing arrow heads indicate bands not apparent in the wild-type WH6 samples. Strains were WH6 (WT WH6), WH6-21G (Δ*gvgR*), WH6-23G (Δ*gvgA*), WH6-24G (Δ*gvgB*), WH6-25G (Δ*gvgC*), WH6-26G (Δ*gvgD*), and WH6-27G (Δ*gvgE*), WH6-28G (Δ*gvgF*), WH6-29G (Δ*gvgG*), WH6-30G (Δ*gvgH*), WH6-31G (Δ*gvgI*), WH6-32G (Δ*gvgJ*), and WH6-33G (Δ*gvgK*).

**Table 1.** Anti-*Erwinia amylovora* activity of WH6 mutant and complemented strains in the agar diffusion assay*

| Genotype | Strain | Anti-*E. amylovora* Activity† |  |  |
| --- | --- | --- | --- | --- |
|  |  | Mutant | Complement |  |
| $\Delta gvgR$ | WH6-21G | $0.0 \pm 0.0$ | pNBW5248N2 | $12.3 \pm 0.13$ |
| $\Delta gvgA$ | WH6-23G | $0.0 \pm 0.0$ | pEVW5249N2 | $11.5 \pm 0.09$ |
| $\Delta gvgB$ | WH6-24G | $0.0 \pm 0.0$ | pEVW5250 | $14.0 \pm 0.19$ |
| $\Delta gvgC$ | WH6-25G | $0.0 \pm 0.0$ | pEVW5251 | $16.2 \pm 0.24$ |
| $\Delta gvgD$ | WH6-26G | $18.1 \pm 0.25$ | N/A | N/A |
| $\Delta gvgE$ | WH6-27G | $14.7 \pm 0.04$ | N/A | N/A |
| $\Delta gvgF$ | WH6-28G | $0.0 \pm 0.0$ | pEVW5254 | $16.7 \pm 0.06$ |
| $\Delta gvgG$ | WH6-29G | $0.0 \pm 0.0$ | pEVW5255 | $16.2 \pm 0.17$ |
| $\Delta gvgH$ | WH6-30G | $0.0 \pm 0.0$ | pEVW5256 | $20.3 \pm 0.25$ |
| $\Delta gvgI$ | WH6-31G | $18.4 \pm 0.34$ | pEVW5257 | $19.1 \pm 0.34$ |
| $\Delta gvgJ$ | WH6-32G | $6.2 \pm 0.21$ | pEVW5258 | $14.1 \pm 0.22$ |
| $\Delta gvgK$ | WH6-33G | $17.4 \pm 0.20$ | pEVW5259 | $14.3 \pm 0.14$ |

\* Area of zone of inhibition in cm<sup>2</sup> shown as the average of three replicates ± SE.
† A typical zone of inhibition for wild-type *P. fluorescens* WH6 is $18.6 \pm 0.45$ .

Mutations in the remaining three genes, *gvgI, gvgJ,* and *gvgK*, led to more complex phenotypes. When *gvgI*, encoding a formyltransferase, was deleted, the filtrate behaved similarly to wild type in the agar diffusion assay (Table 1). However, the activity in the germination arrest assay was reduced (Fig. 3) and the TLC chromatograms were markedly different (Fig. 4). TLC chromatograms of the extracted *gvgI* mutant filtrates lacked the FVG-specific band. Instead, a distinct bright-blue band appeared in chromatograms stained with ninhydrin (Fig. 4). When the Δ*gvgI* mutation was complemented with the wild-type allele, the TLC profile characteristic of FVG (Fig. 5 (a) and (b)) and the germination arrest activity (Fig. 3) were both restored. Thus, it appears that the formyltransferase mutant produces a bioactive compound that is not FVG. Several attempts to identify this molecule failed as the purification process abolished its biological activity (data not shown).

**Fig 5.**
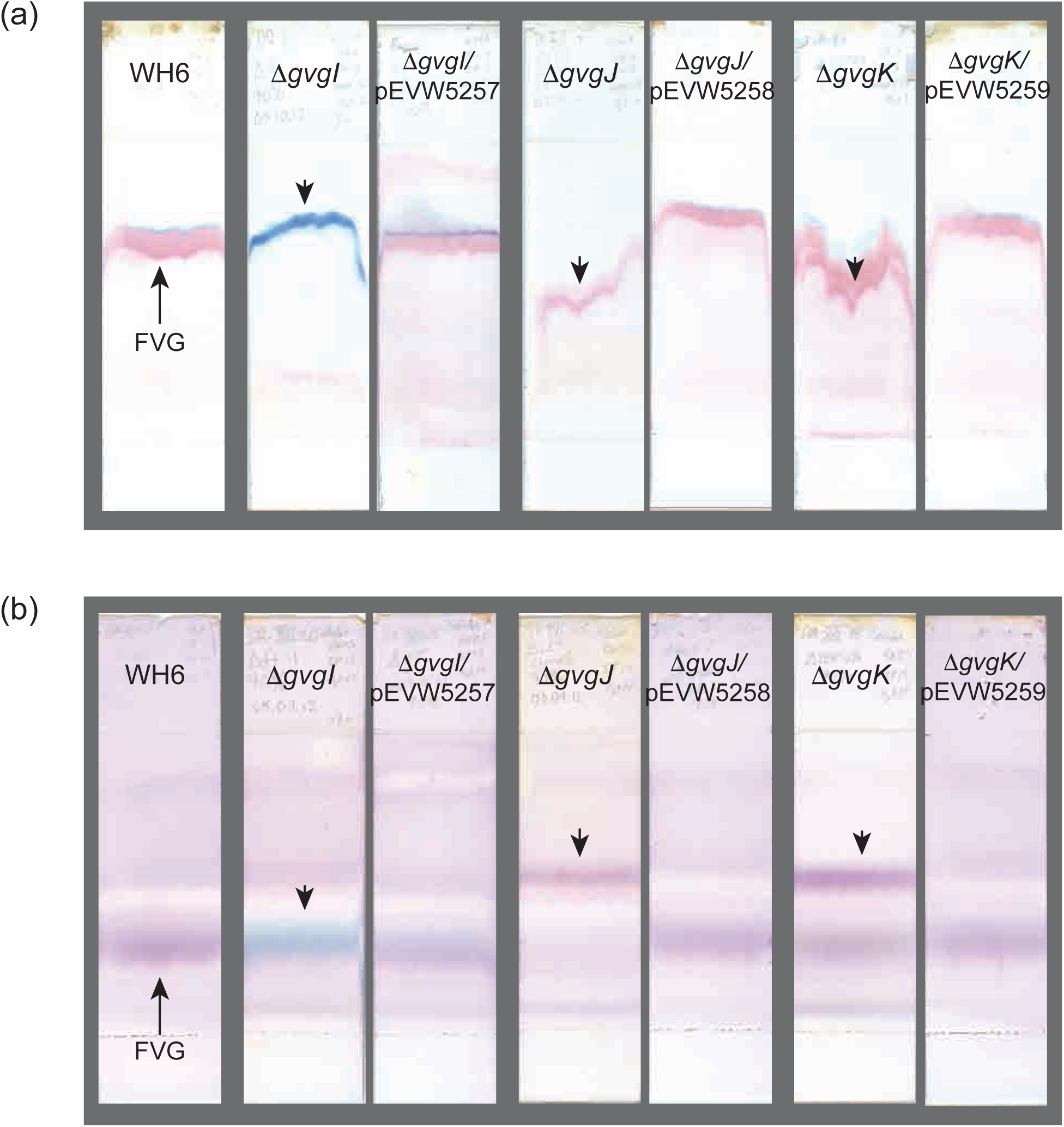
TLC analyses of culture filtrates from *P. fluorescens* WH6 mutant and complemented mutant strains. Ethanol extracts of evaporated culture filtrates were separated by TLC. Strains were WH6-31G (Δ*gvgI*) and its complement, WH6-32G (Δ*gvgJ*) and its complement, and WH6-33G (Δ*gvgK*) and its complement. Downward-pointing arrow heads indicate bands not apparent in the wild-type WH6 samples. (a) Ninhydrin-stained silica TLC plates. (b) Ninhydrin-stained cellulose TLC plates.

The genes *gvgJ* and *gvgK* are both predicted to encode transport proteins in the LysE superfamily (Vrljic *et al.*, 1999) that may facilitate the extracellular export of FVG. Mutant strains with deletions in *gvgJ* or *gvgK* produced distinct phenotypes, however, neither of these genes are absolutely required for FVG production. A mutant strain with a deletion in *gvgJ* displayed reduced activity in the bioassays (Fig. 3, Table 1), indicative of an intermediate phenotype, while mutation of *gvgK* resulted in seemingly wild-type FVG activity in the bioassays (Fig. 3, Table 1). Regardless of their biological activity, silica plate TLC chromatograms of the extracted Δ*gvgJ* or Δ*gvgK* mutant filtrates are quite different from wild-type filtrates (Fig. 4). The blue band typically observed above the FVG band is absent in the Δ*gvgJ* mutant strain and there appears to be an extra pink band in the Δ*gvgK* (Fig. 4) mutant strain. Cellulose TLC chromatograms were subsequently examined to resolve these differences. In the cellulose chromatograms, the bands indicative of FVG appear to be retained in both strains, though at trace levels in the Δ*gvgJ* (Fig. 5 (b)) mutant strain. However, in both strains there is an additional band in the cellulose chromatograms that does not appear in wild-type samples (Fig. 5 (b)). The complementation of the Δ*gvgJ* and Δ*gvgK* mutations with their wild-type alleles restored the FVG phenotype (Fig. 3, Table 1) and eliminated the presence of additional bands in the TLC (Fig. 5 (a) and (b)).

### Mutation of both transporters

The reduction, but not elimination, of FVG with the deletion of *gvgJ* and the similar additional bands in the TLC with the deletion of *gvgJ* or *gvgK* suggested that the two transporters may overlap in function. If GvgJ and GvgK are each able to export FVG, then a strain with a double Δ*gvgJK* deletion would be expected to display a null FVG phenotype. Our attempts to construct a strain with a Δ*gvgJK* genotype, however, were unsuccessful. Although transconjugants harboring the merodiploid were isolated post-conjugation, counter-selection on sucrose led to recovery of wild-type alleles only. This suggested that the double crossover event likely resulted in a lethal mutation, perhaps due to accumulation of a toxic product of the *gvg* cluster.

An alternative strategy for construction of the *gvgJK* deletion was pursued to confirm that the double *gvgJK* deletion is lethal. **(Fig. 6 (a))** The intracellular accumulation of toxic FVG or FVG-related products was avoided by constructing the Δ*gvgJK* mutation in the Δ*gvgC* deletion strain, which does not produce FVG. The double mutation Δ*gvgJK* was mimicked by complementing the triple mutation in the Δ*gvgCHI* mutant strain with *gvgC* under the control of an Ara-inducible promoter. This inducible construct complements the Δ*gvgC* mutation similarly to the equivalent constitutive construct, as indicated by anti-*Erwinia* activity in the agar diffusion assays **(Fig. S2)**. While this complemented Δ*gvgC* strain grew similarly to wild type in the presence of the inducer, the similarly complemented triple mutant grew only slightly (Fig. 6 (b)). This experiment confirms that the lethality of the Δ*gvgJK* mutation is due to product(s) from the *gvg* cluster, either FVG itself or its related byproducts.

**Fig 6.**
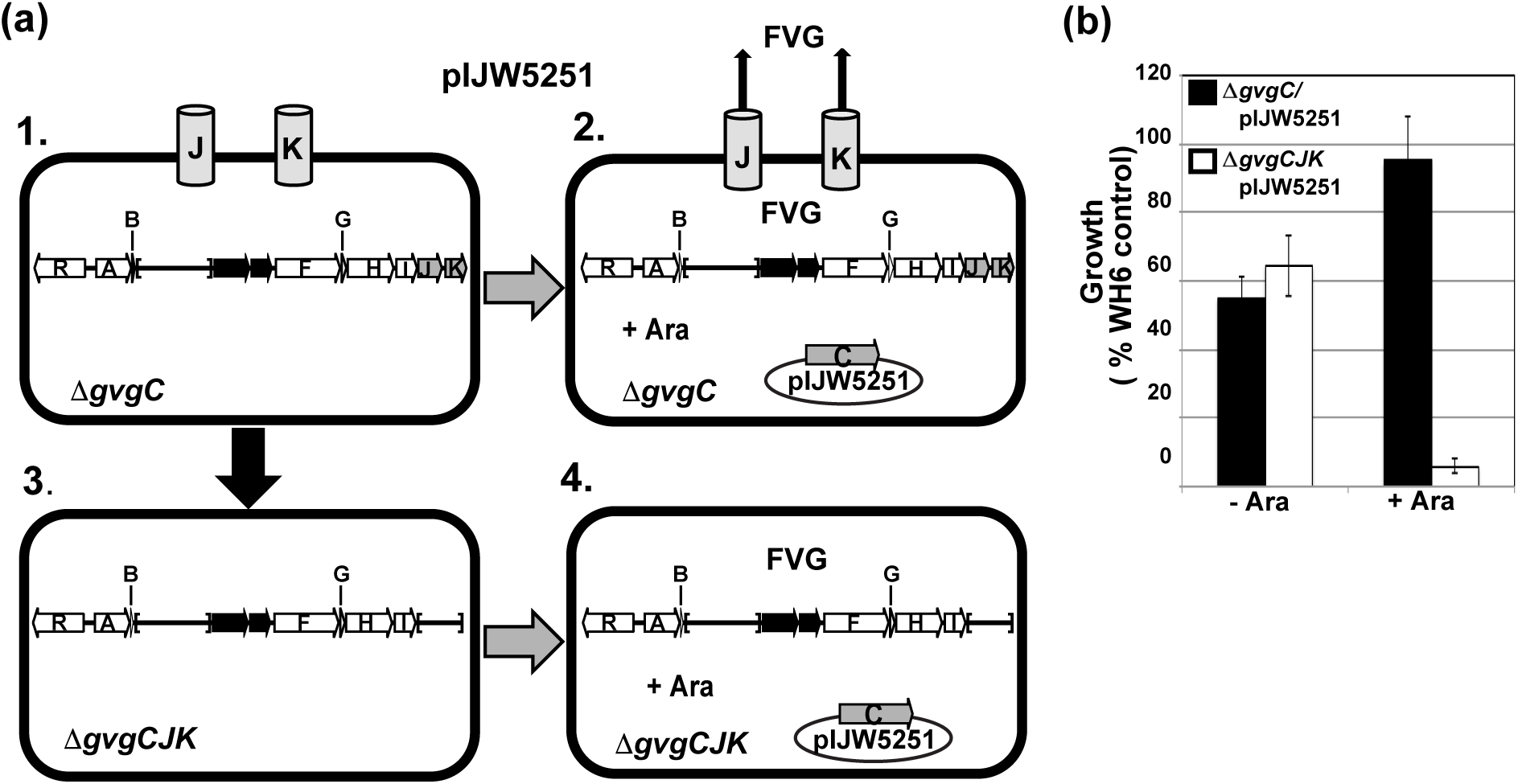
(a) Strategy for deletion of *gvgJ* and *gvgK*. 1. FVG is not produced in ∆*gvgC* mutants. 2. FVG production is restored when ∆*gvgC* is complemented (gray arrow) with pIJW5251 containing *gvgC* under an Ara-inducible promoter. 3. To create the triple mutant, *gvgJ* and *gvgK* are mutated in the ∆*gvgC* background (black arrow). 4. The null-FVG mutant ∆*gvgCJK* is transformed with pIJW5251. In the presence of Ara, this strain should mimic a ∆*gvgJK* mutation, i.e. not viable. (b) Growth of the mutant and complemented mutant strains WH6-25G (∆*gvgC*), WH6-25G/pIJW5251, WH6-34G (∆*gvgCJK),* and WH6-34G/pIJW5251 when in the presence or absence of Ara. Bars represent the average growth of three cultures as a percentage of a wild-type WH6 control ± SD.

### Transcriptional organization of the cluster

As two genes in the center of the cluster are not required for FVG biosynthesis, we were interested in whether the *gvg* cluster is transcribed in a single transcript or in multiple transcripts that bypass these intervening genes. We used several tools to investigate the transcriptional organization of the *gvg* cluster. RT-PCR was performed to determine if the *gvg* genes are coordinately transcribed in a single mRNA transcript. Due to the difficultly in transcribing amplicons larger than 10 kb, we designed the RT-PCR experiment to detect smaller overlapping amplicons over the length of the cluster (Fig. 7 (a)). Overlapping transcripts were detected that span the coding regions of *gvgA* to *gvgK*, but no transcript was detected that extends from *gvgA* into *gvgR* (Fig. 7 (b)). These data suggest that a promoter upstream of *gvgA* drives the expression of the *gvg* cluster in a single transcript including *gvgD* and *gvgE.* This analysis, however, does not eliminate the possibility that smaller regions are also transcribed separately.

**Fig 7.**
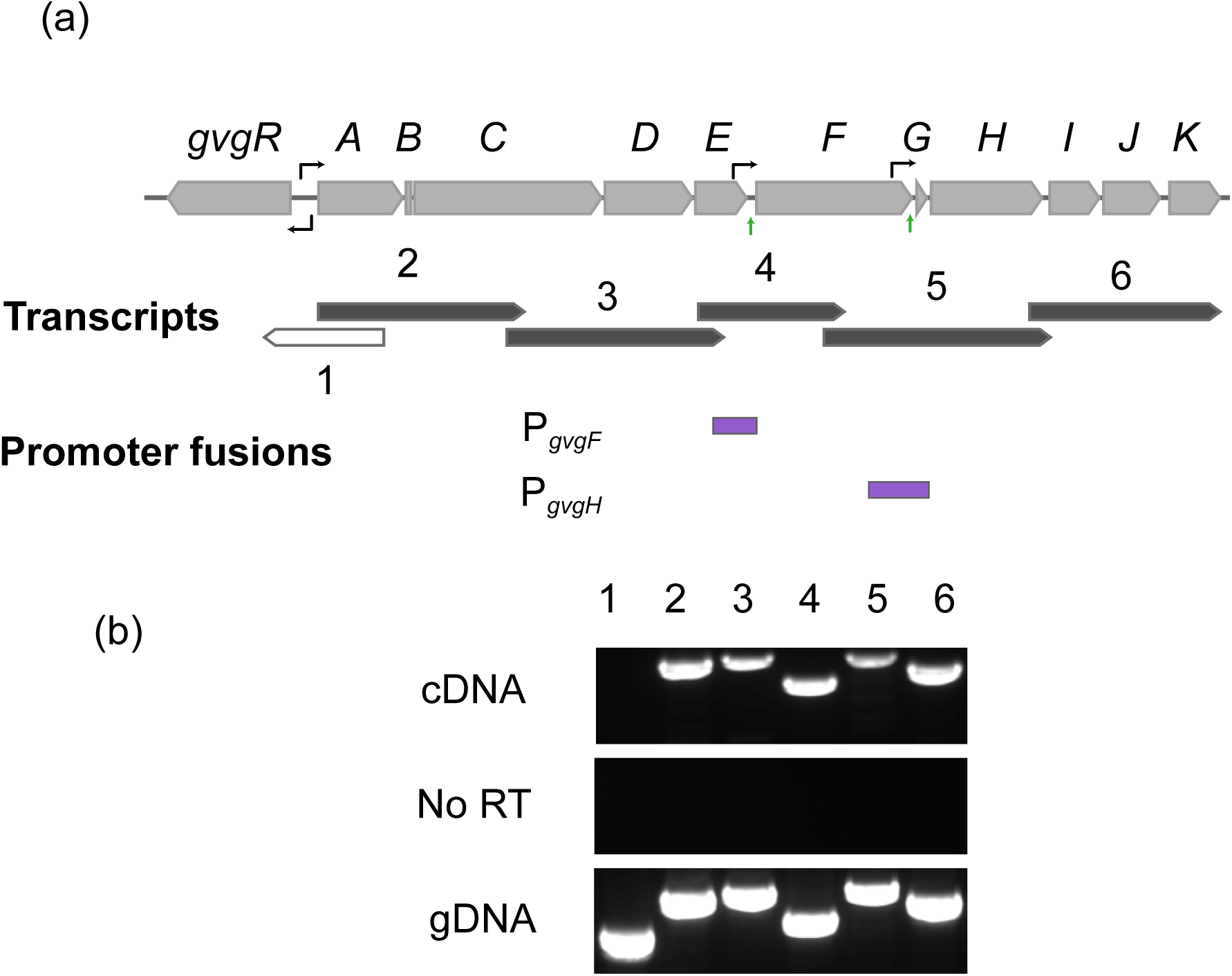
(a) Gene diagram of cluster showing location of overlapping transcripts amplified from *P. fluorescens* WH6 cDNA by RT-PCR. Light gray arrows represent genes in the cluster. The small arrows above the cluster indicate the approximate location of predicted promoters and the small vertical green arrows below the cluster indicate transcriptional start sites determined by 5′ RACE analysis. Solid dark gray arrows represent transcripts amplified from cDNA and the open arrow represents amplification from gDNA (genomic DNA control) but not cDNA. Purple rectangles are sequences cloned into plasmids for expression studies with a *lacZ* reporter. (b) Electrophoresis gel of RT-PCR products amplified from cDNA as well as the negative control with no reverse transcriptase (RT) added and positive control with gDNA as template. Numbers above the lanes correspond to the transcripts in (a).

The possibility of additional transcription from promoters upstream of the CTase-encoding *gvgF* and the aminotransferase-encoding *gvgH* was investigated using 5′ RACE and *lacZ*-promoter fusions. To avoid degraded transcripts, 5′ RACE analysis was performed on mRNA samples enriched for full length transcripts, as described in the Methods section. The longest transcript detected in the *gvgF* 5′ RACE analysis began 29 nt upstream of the predicted *gvgF* start codon (Fig. 7 (a) **and Fig. S3)**. The longest transcript for *gvgH* began 203 nt upstream of the *gvgH* start in the intergenic region between the *gvgF* and *gvgG* (Fig. 7 (a) **and Fig. S3)**. The presence of functional promoters was confirmed by fusing the regions upstream of these genes to a *lacZ* reporter in the wild-type WH6 genetic background. β-galactosidase activity indicative of a functional promoter from the P_*gvgF*_-*lacZ* and P_*gvgH*_-lacZ fusions, along with a P_*lac*_-lacZ control, was qualitatively assessed on LB agar supplemented with X-Gal **(Fig. S4).** A color change in WH6 containing the putative promoter fusions but not in a negative control with *lacZ* only **(Fig. S4)** indicates that the genetic regions upstream of *gvgF* and *gvgH* contain functional promoter sequences. Combined with the RT-PCR and 5′ RACE experiments, these data suggest that the *gvg* gene cluster is transcribed from multiple promoters.

## DISCUSSION

We previously reported the identification of a 13-kb chromosomal region involved in FVG production based on the analysis of Tn*5* insertion libraries in *P. fluorescens* WH6 (Halgren *et al.*, 2013; Kimbrel *et al.*, 2010). This gene cluster is distinct from those involved in the production of other oxyvinylglycines (Cuadrado *et al.*, 2004; Rojas Murcia *et al.*, 2015; Yasuta *et al.*, 2001), which vary considerably despite conserved structural features of the molecules. The objectives of the current study were to: 1) systematically interrogate each gene within the 13-kb region to define the *gvg* cluster, and 2) investigate the transcriptional organization of the gene cluster. Collectively, these data inform our understanding of the biosynthesis of this unusual molecule by providing clues about the enzymatic functions that lead to FVG production. The deletion mutagenesis and functional analysis presented here confirmed the role of genes identified in the Tn*5* screen and also identified six additional genes that are required for FVG biosynthesis.

### Two internal genes are not required for FVG production

The *gvg* gene cluster is disrupted by two genes not required for FVG production. These genes encode an amidinotransferase and a LysE family transporter. The presence of these two genes within the cluster suggests that they may function in combination with other genes in the cluster to produce and export additional compounds. While such a compound has not yet been observed using the standard methodology of detecting FVG, it would likely be hydrophilic and difficult to isolate due to the presence of an amidino group. While unusual, biosynthetic gene clusters that direct the production of multiple compounds have been observed in bacteria. For example, in certain strains of *P. chlororaphis*, phenazine-1-carboxylic acid and a hydroxylated derivative are both produced from the *phz* cluster (Delaney *et al.*, 2001). The two compounds are thought to have different biological roles (Maddula *et al.*, 2008). There are also several examples of superclusters in *Streptomyces* which appear to be merged from smaller clusters (Fischbach, 2009; Mast *et al.*, 2011).

The arrangement of *gvgD* and *gvgE* in the center of the *gvg* cluster is conserved in strains of *Pseudomonas,* such as *P. cannabina* pv. *alisalensis* ES4326 (Halgren *et al.*, 2013). In contrast, *Pantoea ananatis* BRT175 encodes a cluster similar to the *gvg* cluster (Walterson *et al.*, 2014) but lacks the two intervening genes. The BRT175 cluster is responsible for the production of an unidentified compound which, like FVG, inhibits *E. amylovora*. Our results reported here suggest that BRT175 could produce FVG despite the absence of these two genes. Ongoing work in our laboratory is focused on the identification of the active compound from BRT175.

### Small ORFs are required for FVG Production

The requirement of two small ORFs, *gvgB* and *gvgG,* for FVG biosynthesis was surprising. Commonly used annotation pipelines do not, by default, recognize ORFs of this size. Even when annotation pipelines do identify small ORFS, they are often ignored as artifacts (Storz *et al.*, 2014). Indeed, the recent re-annotation of the *P. fluorescens* WH6 genome as part of the NCBI RefSeq project eliminated both of these annotations (NZ_CM001025). However, our complementation data indicate that these small ORFs act out of context of their adjacent genes, therefore they likely encode peptides that are important for FVG production.

There is little known about the role of small peptides in the biosynthesis of secondary metabolites. Bioinformatic analyses of the sequences of GvgB and GvgC do provide some clues as to their function. The DNA sequence of *gvgB* is AT-rich (37% GC content compared to 61% for the entire cluster) and its corresponding amino acid sequence contains an unusually high proportion of lysine residues (8 of 27 total amino acids). There are no predicted transmembrane (TM) domains within the peptide encoded by *gvgB*. In contrast, the DNA sequence of *gvgG* has a GC content (55%) that is similar to the entire *gvg* cluster. The putative peptide encoded by *gvgG* is predicted to consist of 47 amino acids with an alpha-helical TM domain (amino acids 5–24). TM domains are frequently found in proteins encoded by small ORFs. For example, a global characterization found that over 65% of *E. coli* proteins with fewer than 50 amino acids contain a predicted TM domain (Fontaine *et al.*, 2011). Small proteins with TM domains include membrane components that alter membrane characteristics and factors that stabilize large protein complexes (Hobbs *et al.*, 2011).

Small ORFs such as *gvgG* are frequently co-located with an adjacent CTase-encoding gene like *gvgF* (data not shown). Of the CTases that have been functionally characterized, SxtI of the cyanobacterial saxitoxin cluster (Kellmann *et al.*, 2008; Mihali *et al.*, 2009) is the most similar to GvgF (with 55% identity). In *Cylindrospermopsis raciborskii* T3 and other strains of cyanobacteria, two annotated small ORFs are located immediately downstream of the CTase gene. These adjacent genes are both predicted to encode TM domains and are thought to function in combination with the SxtI protein (Kellmann *et al.*, 2008; Mihali *et al.*, 2009). Thus, there may be a common mechanism in which TM-containing small proteins facilitate the function of carbamoyltransferases.

### The role of multiple LysE transporters

Secondary metabolite gene clusters in *Pseudomonas* typically contain a single LysE superfamily transporter (Braun *et al.*, 2010; Brodhagen *et al.*, 2005; Buell *et al.*, 2003; Lee *et al.*, 2010). The *gvg* gene cluster in WH6 breaks this mold by encoding two functional, and somewhat redundant transporters. Based on the presence of specificity-conferring residues reported by Aleshin *et al.* (1999), the LysE superfamily transporters are members of the RhtB family (data not shown), one of three families within the LysE superfamily (Saier, 2000). Although the adjacent positions of *gvgJ* and *gvgK* might suggest they arise from a recent gene duplication event and be similar in sequence, they are actually quite divergent (22% amino acid identity).

Given the phenotype of the mutant strains, it seems likely that GvgJ is the primary exporter of FVG out of the cell, but that GvgK also has some affinity for FVG. When either *gvgJ* or *gvgK* are deleted, a compound accumulates that is not present in TLC chromatograms from wild-type filtrates. This compound could be a precursor or byproduct of FVG biosynthesis. The double Δ*gvgJK* mutation is lethal, suggesting that GvgJ and GvgK can compensate for each other to some extent when only one is deleted. The absence of both transporters, however, seems to result in a toxic accumulation of FVG or other products of the *gvg* cluster within the cell.

### Transcriptional analysis

The *gvg* cluster from *gvgA* to *gvgK* appears to be transcribed in a single transcript which includes *gvgD* and *gvgE*, consistent with a predicted promoter upstream of *gvgA* and a termination signal downstream of *gvgK*. However, 5′ RACE analysis and *lacZ* fusions suggest that there are smaller transcriptional units as well. Multiple modes of transcription may be used to fine-tune the regulation of the cluster. We previously observed multiple transcripts for the *prtIR* genes which encode the PrtI/PrtR sigma factor/anti-sigma factor pair in *P. fluorescens* WH6. The two genes were transcribed together and in separate transcripts with *prtR* transcribed from its own promoter (Okrent *et al.*, 2014). The complex regulation of gene clusters under different environmental conditions and by different sigma factors is commonly found in global studies of transcriptional start sites in bacteria i.e. (Cho *et al.*, 2009; Dötsch *et al.*, 2012; Sharma *et al.*, 2010).

### Conclusions

Functional analysis of the *gvg* cluster reveals a more complete picture of the genes required for FVG production than was available previously. The importance of small ORFs and the involvement of multiple LysE transporters as well as the presence of genes within the *gvg* cluster not required for synthesis of FVG reveal some of the complexities of FVG production. This work thus provides a foundation for subsequent studies to elucidate the biosynthetic pathway of this unique molecule and adds to our knowledge of the diverse genes clusters responsible for oxyvinylglycine biosynthesis.

## AKNOWLEDGMENTS

Research in the laboratory of K.T. is supported by ARS Project # 2072-21410-004-00D. This work was supported by the Agriculture and Food Research Initiative Competitive Grant No. 2012-67012-19868 from the USDA National Institute of Food and Agriculture to R.O.

Technical assistance was provided by Donald Chen. The use of trade, firm, or corporation names in this publication is for the information and convenience of the reader. Such use does not constitute an official endorsement or approval by the United States Department of Agriculture or the Agricultural Research Service of any product or service to the exclusion of others that may be suitable.

## ABBREVIATIONS

Ara, L-arabinose; AVG, aminoethoxyvinylglycine; ECF, Extracytoplasmic function; FRT, *flp* recombinase recognition target; FVG, 4-formylaminooxyvinylglycine; GAF, Germination-Arrest Factor); MVG, 4-methoxyvinylglycine; PLP, pyridoxal phosphate; PMS, Pseudomonas Minimal Salts; 5′ RACE, Rapid Amplification of cDNA Ends; WH6, *Pseudomonas fluorescens* WH6

## FIGURE LEGENDS

**Fig. S1.** Diagram of sequence discrepancies between annotated ORFs in the *Pseudomonas fluorescens* WH6 genome (NCBI CM001025, start sites indicated with dark gray arrows) and ORFs utilized for constructing deletion mutations and plasmids for complements (light gray arrows). The translations are shown under the gene diagrams. (a) Comparison of annotated start sites and utilized start sites for *gvgB* and *gvgC*. In the translation for the overlapping region between *gvgA* and the annotated *gvgB*, the first letter is the translation for *gvgA* and the second for *gvgB*. (b) Comparison of annotated start site and the start site used for constructing deletions mutations and complements of *gvgJ*.

**Fig. S2.** Confirmation that pIJW5251 (*gvgC* under an Ara-inducible promoter) complements the ∆*gvgC* mutation. Strains tested in the using agar diffusion assays for anti-Erwinia activity were wild-type WH6, WH6-25G (Δ*gvgC*), WH6-25G/pEVW5251, and WH6-25G/pIJW5251.

**Fig. S3.** Transcriptional start sites determined from longest sequenced transcripts from 5′ RACE analysis for (a) *gvgF,* with reverse primer within *gvgF* and (b) *gvgG-H,* with reverse primer within *gvgH*.

**Fig. S4.** β-galactosidase activity was qualitatively assessed on LB agar supplemented with X-gal for *lacZ* promoter fusions P_*lac*_-lacZ, P_gvgF_-lacZ and *P_gvgH_-lacZ* along with a promoter-less *lacZ* negative control in the *P. fluorescens* WH6 background. Ten clones were tested from each transformation of *P. fluorescens* WH6.

**Table S1.** Bacterial strains used in this study

**Table S2.** Bacterial plasmids used in this study

**Table S3.** Primers used in this study

**Table S4.** Primers for overlap extension PCR to construct deletions and restriction sites for cloning PCR fragments into pEX18-Km or pEX18-Tc vectors

**Table S5.** Primers for amplification of the FRT-*kan*-FRT fragment for cloning into the deletion constructs

**Table S6.** Primers used for confirmation of deletion mutants

**Table S7.** Primers used to make constructs for complementation or *lacZ* reporter fusions

**Table S8.** Primers used in RT-PCR for transcript identification

**Table S9.** Reverse primers for 5' RACE analysis

**Table S10.** Germination arrest activity from filtrates of mutant strains of *P. fluorescens* WH6

**Table S11.** Germination arrest activity from filtrates of complemented mutant strains of *P. fluorescens* WH6

## References

Aleshin, V. V., Zakataeva, N. P. & Livshits, V. A. (1999). A new family of amino-acid-efflux proteins. Trends Biochem Sci 24, 133–135.

Armstrong, D., Azevedo, M., Mills, D., Bailey, B., Russell, B., Groenig, A., Halgren, A., Banowetz, G. & McPhail, K. (2009). Germination-Arrest Factor (GAF): 3. Determination that the herbicidal activity of GAF is associated with a ninhydrin-reactive compound and counteracted by selected amino acids. Biol Control 51, 181–190.

Banowetz, G. M., Azevedo, M. D., Armstrong, D. J., Halgren, A. B. & Mills, D. I. (2008). Germination-Arrest Factor (GAF): Biological properties of a novel, naturally-occurring herbicide produced by selected isolates of rhizosphere bacteria. Biol Control 46, 380–390.

Banowetz, G. M., Azevedo, M. D., Armstrong, D. J. & Mills, D. I. (2009). Germination arrest factor (GAF): Part 2. Physical and chemical properties of a novel, naturally occurring herbicide produced by *Pseudomonas fluorescen*s strain WH6. Biol Control 50, 103–110.

Berkowitz, D. B., Charette, B. D., Karukurichi, K. R. & McFadden, J. M. (2006). α-Vinylic amino acids: occurrence, asymmetric synthesis, and biochemical mechanisms. Tetrahedron: Asymmetry 17, 869–882.

Braun, S. D., Hofmann, J., Wensing, A., Ullrich, M. S., Weingart, H., Völksch, B. & Spiteller, D. (2010). Identification of the biosynthetic gene cluster for 3-methylarginine, a toxin produced by *Pseudomonas syringae* pv. *syringae* 22d/93. Appl Environ Microbiol 76, 2500–2508.

Brodhagen, M., Paulsen, I. & Loper, J. E. (2005). Reciprocal regulation of pyoluteorin production with membrane transporter gene expression in *Pseudomonas fluorescens* Pf-5. Appl Environ Microbiol 71, 6900–6909.

Buell, C. R., Joardar, V., Lindeberg, M., Selengut, J., Paulsen, I. T., Gwinn, M. L., Dodson, R. J., Deboy, R. T., Durkin, A. S. & other authors (2003). The complete genome sequence of the *Arabidopsis* and tomato pathogen *Pseudomonas syringae* pv. *tomato* DC3000. Proc Natl Acad Sci USA 100, 10181–10186.

Çetinbaş, M., Butar, S., Onursal, C. E. & Koyuncu, M. A. (2012). The effects of pre-harvest ReTain [aminoethoxyvinylglycine (AVG)] application on quality change of 'Monroe' peach during normal and controlled atmosphere storage. Sci Hortic 147, 1–7.

Cho, B.-K., Zengler, K., Qiu, Y., Park, Y. S., Knight, E. M., Barrett, C. L., Gao, Y. & Palsson, B. O. (2009). The transcription unit architecture of the *Escherichia coli* genome. Nat Biotech 27, 1043–1049.

Choi, K.-H., Kumar, A. & Schweizer, H. P. (2006). A 10-min method for preparation of highly electrocompetent *Pseudomonas aeruginosa* cells: application for DNA fragment transfer between chromosomes and plasmid transformation. J Microbiol Methods 64, 391–397.

Choi, K.-H., Mima, T., Casart, Y., Rholl, D., Kumar, A., Beacham, I. R. & Schweizer, H. P. (2008). Genetic tools for select-agent-compliant manipulation of *Burkholderia pseudomallei*. Appl Environ Microbiol 74, 1064–1075.

Cuadrado, Y., Fernández, M., Recio, E., Aparicio, J. F. & Martín, J. F. (2004). Characterization of the *ask-asd* operon in aminoethoxyvinylglycine-producing *Streptomyces* sp. NRRL 5331. Appl Microbiol Biotechnol 64, 228–236.

Datsenko, K. A. & Wanner, B. L. (2000). One-step inactivation of chromosomal genes in *Escherichia coli* K-12 using PCR products. Proc Natl Acad Sci USA 97, 6640–6645.

Delaney, S. M., Mavrodi, D. V., Bonsall, R. F. & Thomashow, L. S. (2001). phzO, a gene for biosynthesis of 2-hydroxylated phenazine compounds in *Pseudomonas aureofaciens* 30-84. J Bacteriol 183, 318–327.

Dötsch, A., Eckweiler, D., Schniederjans, M., Zimmermann, A., Jensen, V., Scharfe, M., Geffers, R. & Häussler, S. (2012). The *Pseudomonas aeruginosa* transcriptome in planktonic cultures and static biofilms using RNA sequencing. PLoS ONE 7, e31092.

Elliott, L., Azevedo, M., Mueller-Warrant, G. & Horwath, W. (1998). Weed control with rhizobacteria. Soil Sci Agrochem Ecol 33, 3–7.

Figurski, D. H. & Helinski, D. R. (1979). Replication of an origin-containing derivative of plasmid RK2 dependent on a plasmid function provided in *trans*. Proc Natl Acad Sci USA 76, 1648–1652.

Fischbach, M. A. (2009). Antibiotics from microbes: converging to kill. Curr Opin Microbiol 12, 520–527.

Fontaine, F., Fuchs, R. T. & Storz, G. (2011). Membrane localization of small proteins in *Escherichia coli*. J Biol Chem 286, 32464–32474.

Halgren, A., Azevedo, M., Mills, D., Armstrong, D., Thimmaiah, M., McPhail, K. & Banowetz, G. (2011). Selective inhibition of *Erwinia amylovora* by the herbicidally active germination-arrest factor (GAF) produced by *Pseudomonas* bacteria. J Appl Microbiol 111, 949–959.

Halgren, A., Maselko, M., Azevedo, M., Mills, D., Armstrong, D. & Banowetz, G. (2013). Genetics of germination-arrest factor (GAF) production by *Pseudomonas fluorescens* WH6: identification of a gene cluster essential for GAF biosynthesis. Microbiology 159, 36–45.

Hobbs, E. C., Fontaine, F., Yin, X. & Storz, G. (2011). An expanding universe of small proteins. Curr Opin Microbiol 14, 167–173.

House, B. L., Mortimer, M. W. & Kahn, M. L. (2004). New recombination methods for *Sinorhizobium meliloti* genetics. Appl Environ Microbiol 70, 2806–2815.

Kellmann, R., Mihali, T. K., Jeon, Y. J., Pickford, R., Pomati, F. & Neilan, B. A. (2008). Biosynthetic intermediate analysis and functional homology reveal a saxitoxin gene cluster in cyanobacteria. Appl Environ Microbiol 74, 4044–4053.

Kimbrel, J. A., Givan, S. A., Halgren, A. B., Creason, A. L., Mills, D. I., Banowetz, G. M., Armstrong, D. J. & Chang, J. H. (2010). An improved, high-quality draft genome sequence of the Germination-Arrest Factor-producing *Pseudomonas fluorescens* WH6. BMC Genomics 11, 522.

Kovach, M. E., Elzer, P. H., Hill, D. S., Robertson, G. T., Farris, M. A., Roop, R. M. & Peterson, K. M. (1995). Four new derivatives of the broad-host-range cloning vector pBBR1MCS, carrying different antibiotic-resistance cassettes. Gene 166, 175–176.

Lee, X., Fox, Á., Sufrin, J., Henry, H., Majcherczyk, P., Haas, D. & Reimmann, C. (2010). Identification of the biosynthetic gene cluster for the *Pseudomonas aeruginosa* antimetabolite L-2-amino-4-methoxy-*trans*-3-butenoic acid. J Bacteriol 192, 4251–4255.

Lee, X., Azevedo, M. D., Armstrong, D. J., Banowetz, G. M. & Reimmann, C. (2013). The *Pseudomonas aeruginosa* antimetabolite L-2-amino-4-methoxy-trans-3-butenoic acid inhibits growth of *Erwinia amylovora* and acts as a seed germination-arrest factor. Environ Microbiol Rep 5, 83–89.

Liu, Y., Rainey, P. B. & Zhang, X.-X. (2014). Mini-Tn7 vectors for studying post-transcriptional gene expression in *Pseudomonas*. J Microbiol Methods 107, 182–185.

Maddula, V. S. R. K., Pierson, E. A. & Pierson, L. S. (2008). Altering the ratio of phenazines in *Pseudomonas chlororaphis (aureofaciens)* strain 30-84: Effects on biofilm formation and pathogen inhibition. J Bacteriol 190, 2759–2766.

Markowitz, V. M., Chen, I.-M. A., Palaniappan, K., Chu, K., Szeto, E., Pillay, M., Ratner, A., Huang, J., Woyke, T. & other authors (2014). IMG 4 version of the integrated microbial genomes comparative analysis system. Nucleic Acids Res 42, D560–D567.

Mast, Y., Weber, T., Gölz, M., Ort-Winklbauer, R., Gondran, A., Wohlleben, W. & Schinko, E. (2011). Characterization of the 'pristinamycin supercluster' of *Streptomyces pristinaespiralis*. Microb Biotechnol 4, 192–206.

McPhail, K. L., Armstrong, D. J., Azevedo, M. D., Banowetz, G. M. & Mills, D. I. (2010). 4-Formylaminooxyvinylglycine, an herbicidal germination-arrest factor from *Pseudomonas* rhizosphere bacteria. J Nat Prod 73, 1853–1857.

Mihali, T., Kellmann, R. & Neilan, B. (2009). Characterisation of the paralytic shellfish toxin biosynthesis gene clusters in *Anabaena circinalis* AWQC131C and *Aphanizomenon* sp. NH-5. BMC Biochem 10, 8.

Mitchell, R. E. & Frey, E. J. (1988). Rhizobitoxine and hydroxythreonine production by *Pseudomonas andropogonis* strains, and the implications to plant disease. Physiol Mol Plant Pathol 32, 335–341.

Naville, M., Ghuillot-Gaudeffroy, A., Marchais, A. & Gautheret, D. (2011). ARNold: A web tool for the prediction of Rho-independent transcription terminators. RNA Biol 8, 11–13.

Newman, J. R. & Fuqua, C. (1999). Broad-host-range expression vectors that carry the L-arabinose-inducible *Escherichia coli* araBAD promoter and the *araC* regulator. Gene 227, 197–203.

Okrent, R. A., Halgren, A. B., Azevedo, M. D., Chang, J. H., Mills, D. I., Maselko, M., Armstrong, D. J., Banowetz, G. M. & Trippe, K. M. (2014). Negative regulation of germination-arrest factor production in *Pseudomonas fluorescens* WH6 by a putative extracytoplasmic function sigma factor. Microbiology 160, 2432–2442.

Owens, L. D., Thompson, J. F., Pitcher, R. & Williams, T. (1972). Structure of rhizobitoxine, an antimetabolic enol-ether amino-acid from *Rhizobium japonicum*. J Chem Soc, Chem Commun, 714–714.

Parthier, C., Görlich, S., Jaenecke, F., Breithaupt, C., Bräuer, U., Fandrich, U., Clausnitzer, D., Wehmeier, U. F., Böttcher, C. & other authors (2012). The O-carbamoyltransferase TobZ catalyzes an ancient enzymatic reaction. Angew Chem Int Ed 51, 4046–4052.

Percudani, R. & Peracchi, A. (2003). A genomic overview of pyridoxal-phosphate-dependent enzymes. EMBO Reports 4, 850–854.

Petersen, T. N., Brunak, S., von Heijne, G. & Nielsen, H. (2011). SignalP 4.0: discriminating signal peptides from transmembrane regions. Nat Methods 8, 785–786.

Pruess, D. L., Scannell, J. P., Kellett, M., Ax, H. A., Janecek, J., Williams, T. H., Stempel, A. & Berger, J. (1974). Antimetabolites produced by microorganisms. X. L-2-amino-4-(2-aminoethoxy)-trans-3-butenoic acid. J Antibiot 27, 229–233.

Rojas Murcia, N., Lee, X., Waridel, P., Maspoli, A., Imker, H. J., Chai, T., Walsh, C. T. & Reimmann, C. (2015). The *Pseudomonas aeruginosa* antimetabolite L-2-amino-4-methoxy-trans-3-butenoic acid (AMB) is made from glutamate and two alanine residues via a thiotemplate-linked tripeptide precursor. Front Microbiol 6, 170.

Saier, M. H. (2000). Families of transmembrane transporters selective for amino acids and their derivatives. Microbiology 146, 1775–1795.

Scannell, J. P., Pruess, D., Blount, J. F., Ax, H. A., Kellett, M., Weiss, F., Demny, T. C., Williams, T. H. & Stempel, A. (1975). Antimetabolites produced by microorganisms. XII. (S)-alanyl-3-[alpha-(S)-chloro-3-(S)-hydroxy 2-oxo-3-azetidinylmethyl]-(S)-alanine, a new beta-lactam containing natural product. J Antibiot 28, 1–6.

Sharma, C. M., Hoffmann, S., Darfeuille, F., Reignier, J., Findeisz, S., Sittka, A., Chabas, S., Reiche, K., Hackermuller, J. & other authors (2010). The primary transcriptome of the major human pathogen *Helicobacter pylori*. Nature 464, 250–255.

Solovyev, V. & Salamov, A. (2011). Automatic annotation of microbial genomes and metagenomic sequences. In Metagenomics and its Applications in Agriculture, Biomedicine and Environmental Studies, pp. 61–78. Edited by R. W. Li. Hauppauge, NY: Nova Science Publishers.

Storz, G., Wolf, Y. I. & Ramamurthi, K. S. (2014). Small proteins can no longer be ignored. Annu Rev Biochem 83, 753–777.

Vrljic, M., Garg, J., Bellmann, A., Wachi, S., Freudl, R., Malecki, M. J., Sahm, H., Kozina, V. J., Eggeling, L. & other authors (1999). The LysE superfamily: topology of the lysine exporter LysE of *Corynebacterium glutamicum*, a paradyme for a novel superfamily of transmembrane solute translocators. J Mol Microbiol Biotechnol 1, 327–336.

Walterson, A. M., Smith, D. D. N. & Stavrinides, J. (2014). Identification of a *Pantoea* biosynthetic cluster that directs the synthesis of an antimicrobial natural product. PLOS ONE 9, e96208.

Yasuta, T., Okazaki, S., Mitsui, H., Yuhashi, K. I., Ezura, H. & Minamisawa, K. (2001). DNA sequence and mutational analysis of rhizobitoxine biosynthesis genes in *Bradyrhizobium elkanii*. Appl Environ Microbiol 67, 4999–5009.

Yu, Y.-B., Adams, D. O. & Yang, S. F. (1979). 1-Aminocyclopropanecarboxylate synthase, a key enzyme in ethylene biosynthesis. Arch Biochem Biophys 198, 280–286.

